# Differential response of *Candida* species morphologies and isolates to fluconazole and boric acid

**DOI:** 10.1101/2022.01.09.475582

**Authors:** Ola E. Salama, Aleeza C. Gerstein

## Abstract

*Candida albicans* is the most prevalent cause of vulvovaginal candidiasis (‘yeast infection’) and recurrent vulvovaginal candidiasis, though the incidence of non-albicans yeast species is increasing. The azole fluconazole is the primary antifungal drug used to treat R/VVC yet isolates from some species have intrinsic resistance to fluconazole, and recurrent infection can occur even with fluconazole-susceptible populations. The second-line broad-spectrum antimicrobial drug, boric acid, is an alternative treatment that has been found to successfully treat complicated VVC infections. Far less is known about how boric acid inhibits growth of yeast isolates in different morphologies compared to fluconazole. We found significant differences in drug resistance and drug tolerance (the ability of a subpopulation to grow slowly in high levels of drug) between *C. albicans, C. glabrata*, and *C. parapsilosis* isolates, with the specific relationships dependent on both drug and phenotype. Population-level variation for both susceptibility and tolerance was broader for fluconazole than boric acid in all species. Unlike fluconazole, which neither prevented hyphal formation nor disrupted mature biofilms, boric acid inhibited *C. albicans* hyphal formation and reduced mature biofilm biomass and metabolic activity in all isolates in a dose-dependent manner. Variation in planktonic response did not generally predict biofilm phenotypes. Overall, our findings illustrate that boric acid is broadly effective at inhibiting growth across many isolates and morphologies, which could explain why it is an effective treatment for R/VVC.

## Introduction

Vulvovaginal candidiasis (VVC), a pathological condition of the lower female reproductive tract, affects ∼75% of females at least once in their life (1, 2). Although patients usually respond well to the first-line treatment (typically fluconazole) (3, 4), ∼8% of females globally will experience recurrent VVC, which can have a significant negative impact on quality of life (4, 5). *Candida albicans* has been the primary cause (75-90%) of VVC, yet the frequency of other species, particularly *C. glabrata* followed by *C. parapsilosis*, is increasing (6–8). In light of the transition to rename *C. glabrata* as *Nakaseomyces glabrata*, (9), we use the acronym NAC to refer to “non-albicans clinical” species, rather than “non-albicans *Candida*” species as has been done historically.

Boric acid (BA) intravaginal suppositories are recommended and have found success as a second-line treatment for NAC infections and when recurrence occurs on first-line treatment, typically fluconazole (FLC) (3, 4, 10, 11). Despite this success, BA remains a second-line treatment because its exact mechanism of action is unknown and much less work has evaluated its broad efficacy and possible long-term effects (12, 13). The minimum inhibitory concentration of BA was previously measured in a relatively small number of *C. albicans, C. glabrata*, and *C. parapsilosis* isolates (13, 14), yet unlike FLC (and many other antifungal drugs), there are no standard clinical methods for drug resistance testing, nor have breakpoints been established for BA planktonic resistance. Furthermore, variation among isolates in fungal drug tolerance, the ability of a subpopulation of drug-susceptible isolates to grow slowly at inhibitory drug concentrations (15–18), has largely been ignored, yet recent studies are beginning to implicate tolerance as a factor in predicting drug efficacy (16–18).

Morphological plasticity is an important virulence trait in many contexts for *Candida* species (19). The yeast to hyphal transition is critical for biofilm stability, penetration of host epithelial cells, and escape from host phagocytes (20, 21). Hyphal formation is also important for biofilm formation in *C. albicans* (*C. glabrata* do not form hyphae and *C. parapsilosis* forms pseudohyphae) (22). The involvement of biofilms in RVVC is still being debated (23, 24). However, hyphal forms were recently detected in vaginal lavage fluid from an RVVC patient (25). I*n vivo* mice models showed that *C. albicans* can form biofilms on the vaginal mucosa (23, 24) and genes involved in hyphal morphogenesis biofilm formation were detected in vaginal lavage (25). FLC was shown previously not to inhibit hyphal formation (26). However, low BA concentrations (0.02 mg/mL) can disrupt the cytoskeleton of hyphae by changing the actin distribution from the apical to the isotropic pattern (27), and higher BA concentrations (10 mg/mL or 50 mg/mL) were shown to inhibit hyphal formation in two *C. albicans* isolates (13). BA has also been found to reduce the biomass of mature biofilms relative to biofilms growing without drug (13); it was unclear, however, whether this reflected the drug simply stopping further growth, or whether boric acid acted to reduce the biofilm below pre-drug treatment levels.

To quantify the diversity of BA phenotypic responses among different species, we compared FLC and BA planktonic susceptibility and tolerance among 235 clinical isolates of *C. albicans, C. glabrata* and *C. parapsilosis*. We also quantify the impact of FLC and BA on *C. albicans* yeast to hyphal transition (leading to biofilm formation), and on *C. albicans* mature biofilms. We found significant differences among species for drug resistance and tolerance, and a consistent increase in the variance among isolates for both drug response phenotypes in FLC compared to BA. We also found that unlike FLC, BA is effective at inhibiting the yeast-to-hyphal transition and thus biofilm formation, and can effectively break apart mature biofilms. Combined, this demonstrates multiple pathways where BA is more effective at inhibiting *Candida* species growth compared to fluconazole.

## Results

### Variation in Drug Susceptibility and Tolerance

Fluconazole (FLC) and boric acid (BA) drug susceptibility and tolerance were measured for 235 *Candida* isolates (165 *C. albicans*, 50 *C. glabrata*, 20 *C. parapsilosis*) using *diskImageR*, a computational analysis tool that quantifies drug response from imagers of disk diffusion assays (15). Susceptibility was measured as RAD_20_ (the radius where 20% of growth reduction occurs), while tolerance was measured as FoG_20_ (the fraction of the population that is able to grow above RAD_20_ after 48 h). *C. albicans* isolates on average had higher susceptibility than *C. glabrata* but lower susceptibility than *C. parapsilosis*, and higher tolerance than either in FLC (Figure 1, left panels; Kruskal-Wallis rank-sum test; susceptibility: χ^2^ = 63.43 df = 2, *P* < 0.0001, species differences determined by post-hoc pairwise Wilcoxon tests with the (28) *P* adjustment; tolerance: χ^2^ = 82.04, df = 2, *P* < 0.0001). In BA, *C. albicans* isolates had higher susceptibility than both NAC species, and lower tolerance than *C. glabrata* (susceptibility: χ^2^ = 39.8, df = 2, *P* < 0.0001; tolerance: χ^2^ = 97.9, df = 2, *P* < 0.0001). The magnitude of isolate variation in BA was significantly less than in FLC for both *C. albicans* and *C. glabrata*, the two species with a sufficiently large sample size (Fligner-Killeen test of homogeneity; susceptibility: *C. albicans*, χ^2^ = 23.4, df = 1, *P* < 0.0001; *C. glabrata*, χ^2^ = 7.8, df = 1, *P* = 0.005; tolerance: *C. albicans*, χ^2^ = 116.59, df = 1, *P* < 0.0001; *C. glabrata*, χ^2^ = 4.94, df = 1, *P* = 0.03).

**Figure 1.**
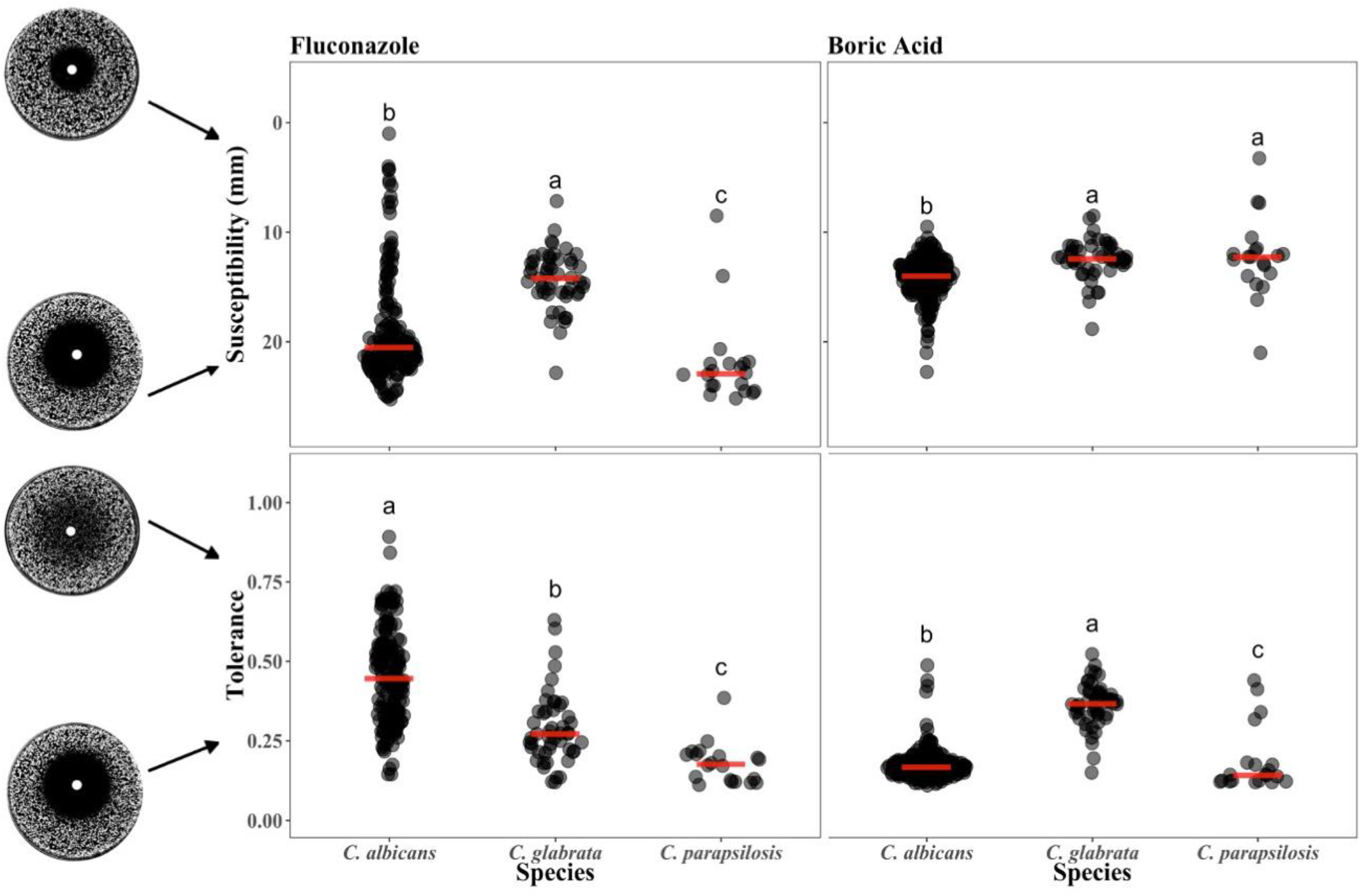
Susceptibility and tolerance for FLC and BA was quantified for 165 *C. albicans*, 50 *C. glabrata*, and 20 *C. parapsilosis* isolates. Note that susceptibility is measured as RAD_20_ and the y-axis is reversed so that the less susceptible (more resistant) isolates are towards the top. Tolerance is measured FoG_20_ from *diskImageR* analysis. Letters indicate the statistical differences among-species in each panel from a post-hoc pairwise Wilcox test with the Benjamini and Hochberg (1995) p-value adjustment; when species do not share a letter, they are significantly different from each other (*P* < 0.05).

There was a significant positive correlation for FLC susceptibility and tolerance within isolates of the same species: isolates that had lower susceptibility (i.e., higher resistance levels) also tended to have higher tolerance (Figure 2A, Spearman’s rank correlation, for *C. albicans:* rho = −0.197, S = 571000, *P* = 0.02 *C. glabrata*: rho = −0.49, S = 29249, *P* < 0.0001, *C. parapsilosis*: rho = −0.73, S = 2295, *P* < 0.0001). In BA, the correlation was significant for *C. glabrata* and *C. parapsilosis* (Figure 2B, *C. glabrata*, rho = −0.42, S = 27802, *P* = 0.003; *C. parapsilosis*, rho = −0.63, S = 2173, *P* = 0.003) but not *C. albicans* (rho = 0.07, S = 444639 *P* = 0.420). There was no correlation in susceptibility and tolerance between drugs, indicating that the mode of action for BA differs from that of FLC, as the isolates that are less susceptible or more tolerant in one drug do not tend to have improved growth in the other (Figure 2C&D, Spearman’s rank correlation, susceptibility, *C. albicans*: S = 500605, *P* = 0.56; *C. glabrata*: S = 19549, *P* = 0.99; *C. parapsilosis:* S = 975, *P* = 0.26; tolerance, *C. albicans:* S = 490557, *P* = 0.74; *C. glabrata*: S = 15401, *P* = 0.14; *C. parapsilosis:* S = 1332, *P* = 0.995).

**Figure 2.**
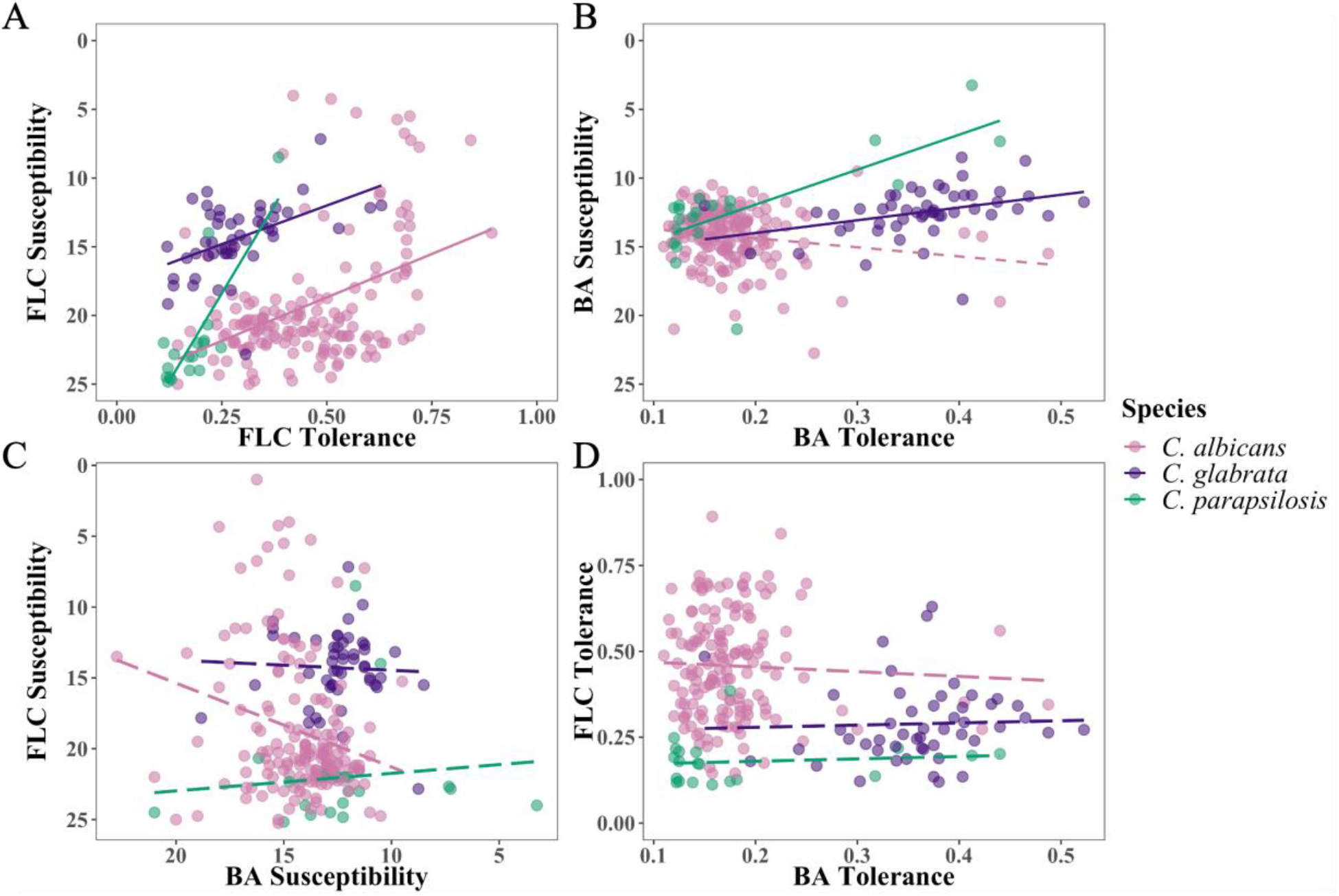
Association within and between drug response parameter and drugs. (A) susceptibility and tolerance in FLC, (B) susceptibility and tolerance in BA, (C) susceptibility in FLC and BA, (D) tolerance in FLC and BA. Solid lines indicate a significant correlation (*P* < 0.05) from a Spearman’s rank correlation test.

### BA Inhibits *C. albicans* Hyphal Formation

We used time-lapse microscopy to examine the ability of eight phylogenetically diverse *C. albicans* isolates to form hyphae and begin biofilm formation in the presence of drug (Figure 3). Visual inspection of images indicated it took only 1-2 hours for all isolates to form hyphae regardless of the FLC concentration (Figure 3B). By contrast, BA affected hyphal formation in a highly dose-dependent manner. In low levels of BA (0.4 and 0.8 mg/mL), there was very little variation in the time to first hyphal formation among isolates. As the level of BA increased, the time to hyphal formation as well as the variation among isolates increased. No hyphae were observed by 24 h at the highest level of BA.

**Figure 3.**
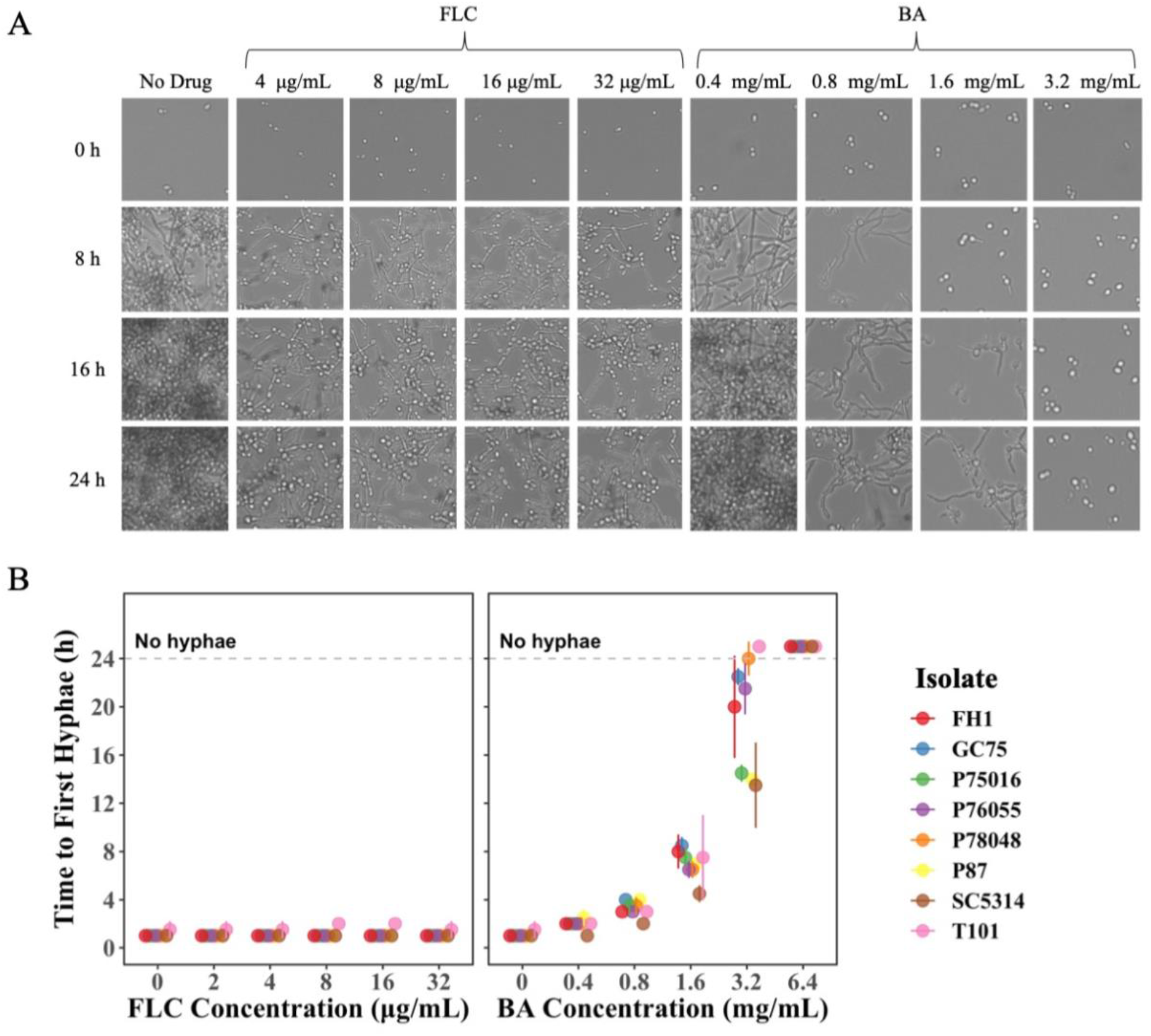
Biofilm formation drug responses of eight *C. albicans* isolates. Fungal cells were initially cultured in liquid YPD at 30 °C, washed and standardized to OD_600_ of 0.01 in RPMI. Cells were inoculated into various concentrations of FLC or BA. The plate was incubated at 37 °C and (A) manually taken out every 1 h for 24 h for scan using Evos FL Auto 2 inverted microscopy. (B) Time to the first hyphal formation was measured by manually going through the images and identifying the hour where the first hypha was observed. Values presented for each isolate are the mean of two biological replicates ± SD.

To further quantify biofilm formation from the time-lapse images, we used a computational pipeline that we recently developed that uses machine learning in the Orbit Image Analysis program (29) and custom R scripts. The pipeline computationally quantifies the percent area covered by cells in each image, and uses this to calculate the biofilm growth rate, the time to reach the growth asymptote (i.e., growth plateau), and percent area covered at the asymptote. At the highest FLC concentration, populations retained 50-100 % of percent area covered by cells at the asymptote relative to no drug (Figure 4A) and had a reduced but still relatively high growth rate (Figure 4B). There was not a clear trend among isolates in the time required to reach the asymptote (Figure 4C). Consistent with the identified variation by eye in the time to hyphal formation, BA reduced the percent area covered by cells at the growth asymptote, decreased the growth rate and increased the time to reach the growth asymptote in a dose-dependent manner (Figure 4A-C). Interestingly, there was more variation in percent area at the growth asymptote, growth rate, and time to reach the growth asymptote among isolates in FLC compared to BA. There was no correlation between FLC drug responses and BA drug responses (Spearman’s rank correlation, percent area covered by cells: S = 1399899, *P* = 0.48; time to reach the growth asymptote S = 1346887, *P* = 0.89; growth rate: S = 1275866, *P* = 0.5447). Overall, unlike FLC, BA effectively inhibits hyphal formation and biofilm formation in a dose-dependent manner.

**Figure 4.**
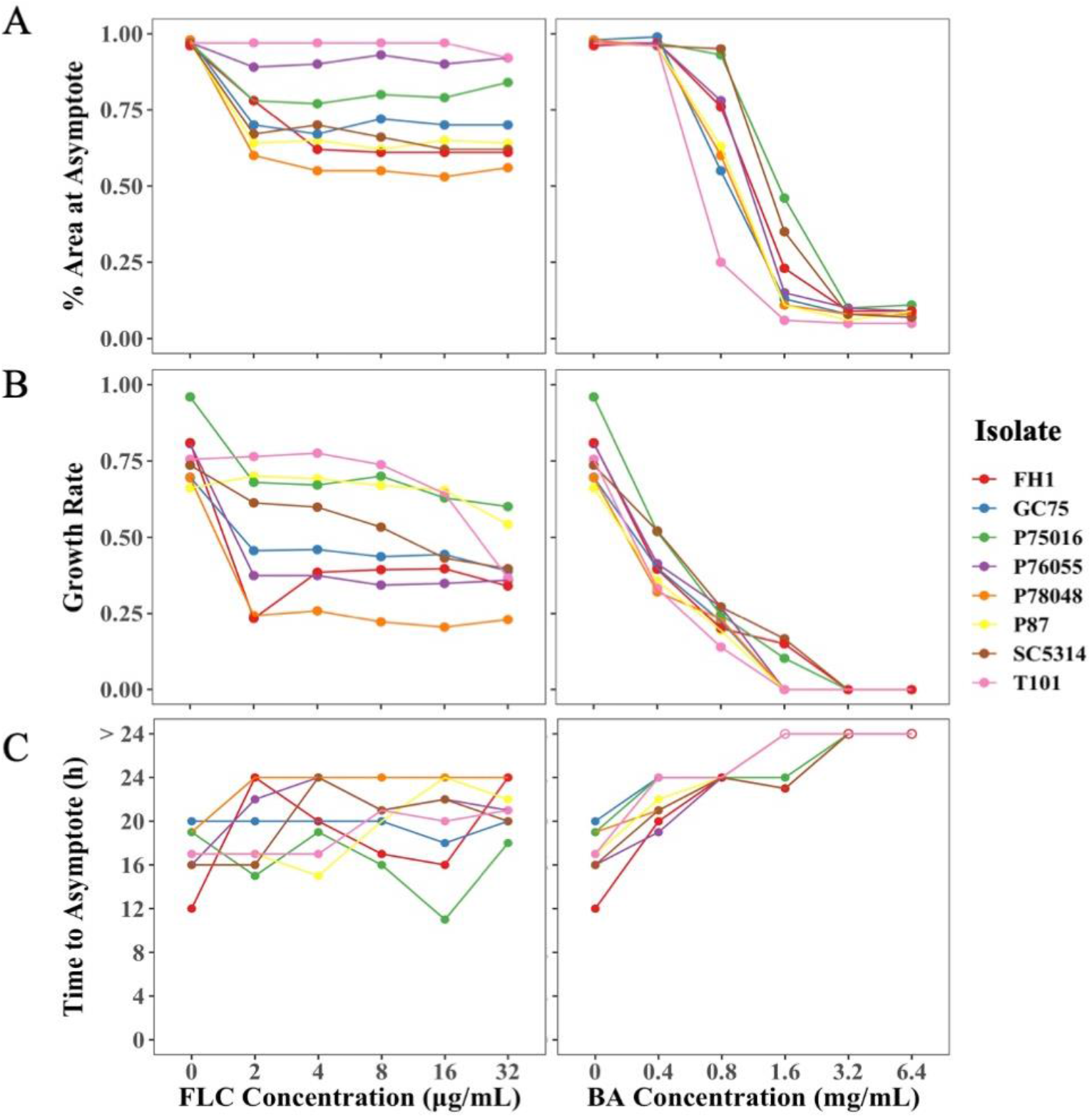
Quantitative image analysis of time lapse images of eight *C. albicans* isolates. Orbit Image Analysis was used to calculate the % area covered by cells for all the images and custom R scripts was used to calculate % area at the asymptote, growth rate and time to reach the asymptote. BA affected biofilm formation in a dose dependent manner; (A) it reduced % area at the asymptote, (B) decreased the growth rate, and (C) increased time to reach the asymptote unlike FLC. Values presented are the mean of two biological replicates.

### BA Eradicates *C. albicans* Mature Biofilms

The ability of drugs to break apart mature *C. albicans* biofilms formed in absence of drug was evaluated by quantifying both biomass and activity. As previously shown by others (24, 30, 31), mature biofilms were largely impervious to FLC; post-drug biomass was higher than the measured pre-drug biomass for all levels of FLC (Figure 5; linear mixed-effect model implemented in the lmer R package (32), with change in biomass as the response variable, level of the drug as the predictor and isolate as a random effect, p-value obtained through the analysis of variance test with Satterthwaite’s method for degrees of freedom; F_1,342_ = 2.98, *P* = 0.09). There was a negative association between FLC concentration and biofilm activity (linear mixed-effect model with change in activity as the response variable, level of the drug as the predictor, and isolate as a random effect, F_1,342_ = 32.46, *P* < 0.0001), driven by activity at the highest level of FLC (model with FLC 32 μg/mL removed from the dataset: F_1,285_ = 0.0937, *P* = 0.76).

**Figure 5.**
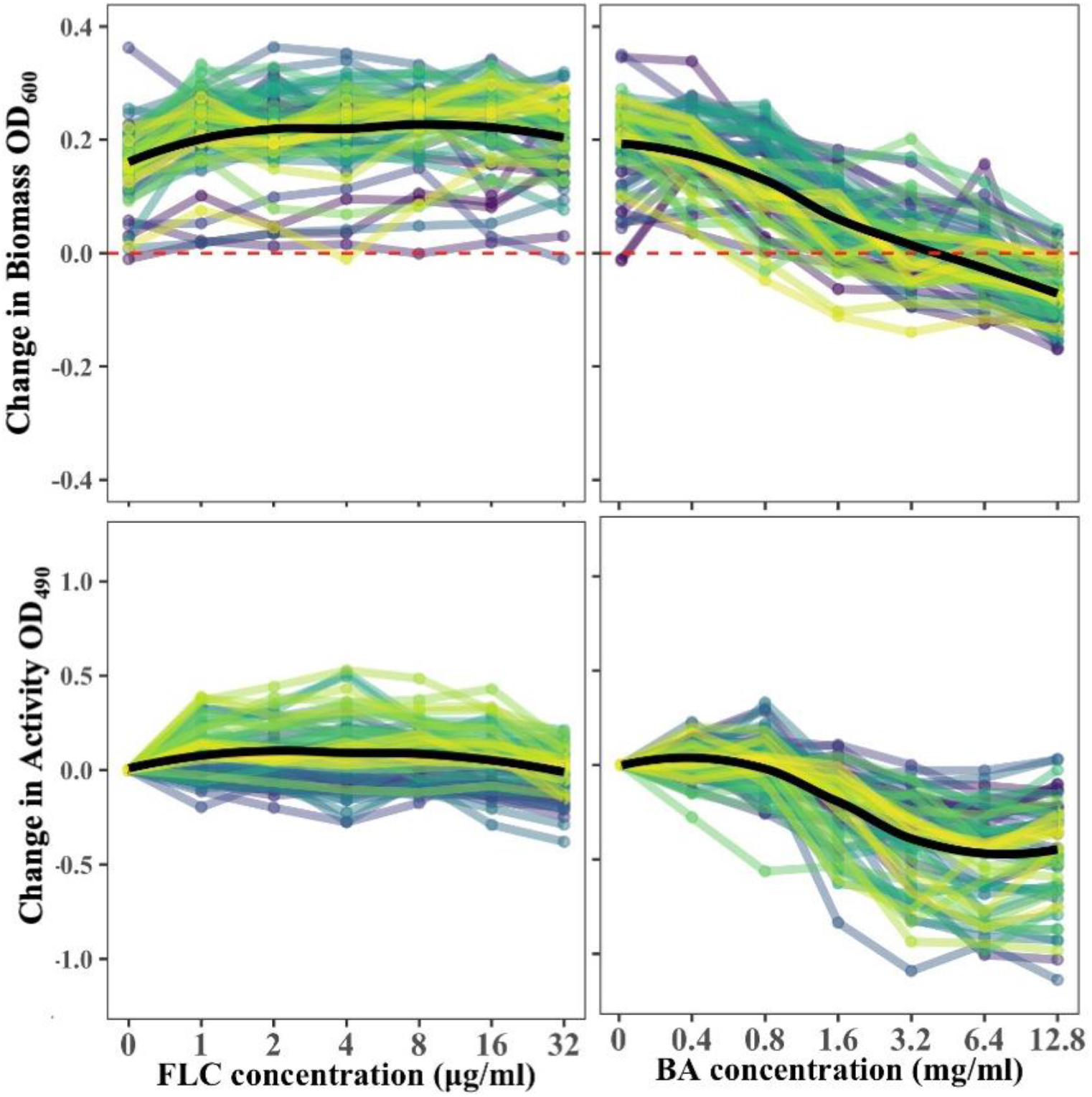
Biofilms drug responses of 63 *C. albicans* isolates. In all panels, the black line indicates the mean across all isolates at each concentration. The line colors are arbitrary and used to visualize different isolates. Biofilms were insensitive to FLC regardless of the FLC concentration. BA eradicated mature biofilms in a dose-dependent manner and reduced activity. Values presented are the means of three biological replicates.

Similar to biofilm formation, BA significantly affected biofilm biomass and activity in a dose-dependent manner (Figure 5; biomass, F_6,342_ = 249.94, *P* < 0.0001; activity, F_6,342_ = 249.94, *P* < 0.0001). Interestingly, the biomass of 48% of isolates and the activity of 57% of isolates increased at 0.4 mg/mL BA relative to no drug, suggesting that BA can stimulate growth at a low concentration. The biofilm biomass at high BA concentrations relative to a biofilm grown without drug was reduced for all isolates. The activity of the remaining biofilm at the highest BA levels for some isolates increased above that of the preceding lower drug level, likely because BA degraded the biofilm matrix allowing XTT to better penetrate the cells.

Planktonic drug responses were partially predictive of biofilm responses, albeit in an inconsistent manner. Isolates with higher planktonic growth tended to have a higher biofilm activity but not higher biomass, while lower susceptibility (higher resistance) did not predict either biofilm biomass or activity (Supplementary Materials Figure 1-4; linear mixed-effect model with the FLC planktonic response as the predictor variables and isolate as a random effect; change in biomass as the response variable: susceptibility, F_1,57_ = 1.59, *P* = 0.21, tolerance, F_1, 54_ = 1.28, *P* = 0.26; change in activity as the response variable, susceptibility, F_1, 57_ = 2.51, *P* = 0.12, tolerance, F_1, 54_ = 5.76, *P* = 0.02). By contrast, reduced BA planktonic susceptibility was associated with higher biofilm activity but not biomass, while tolerance was associated with neither (susceptibility, change in biomass, F_1, 57_ = 1.04, *P* = 0.31, change in activity, F_1, 57_ = 5.55, *P* = 0.02; tolerance, change in biomass, F_1, 57_ = 0.0004, *P* = 0.98; change in activity, F_1, 57_ = 0.07, *P* =0.79). Hence, isolate differences in planktonic drug responses are likely not a good predictor for how a specific isolate will respond to drug when in a biofilm.

## Discussion

We screened isolates of *Candida* species to enhance our understanding of how diverse isolates in multiple morphologies respond to boric acid (BA), an effective drug for treating complicated vulvovaginal candidiasis (VVC). We directly compared responses against fluconazole (FLC), the first-line treatment. Consistent with previous work, compared to *C. albicans*, we found that *C. glabrata* on average had lower susceptibility to both FLC (33, 34) and BA (13, 14) Our work is also in agreement with previous work that found no correlation between FLC and BA susceptibility in 76 *C. albican* isolates (27), implying that the mode of action of FLC is different from that of BA. In both *C. albicans* and *C. glabrata*, we found that the distribution of susceptibility values was broader in FLC than BA. Rosenberg *et al*. (2018) measured susceptibility in 19 *C. albicans* isolates in seven antifungal drugs and similarly found FLC had the broadest distribution (16). Our work suggests BA resistant isolates may be rare as only one *C. parapsilosis* isolate out of 235 *C. albicans, C. glabrata* and *C. parapsilosis* isolates had considerably higher resistance than other isolates. Additional screening with a larger isolate collection is required to establish clinical breakpoints of BA resistance.

We also found significant variation among species and a wider breadth of responses in FLC compared to BA for drug tolerance. There is a growing acknowledgment that fungal drug tolerance may be associated with treatment failure and infection persistence. One study showed that *C. albicans* isolates from patients with persistent candidemia had a higher tolerance to FLC than isolates from patients with non-persistent candidemia (16). Increased tolerance was also found to be a good predictor of FLC efficiency in patients with *C. albicans* bloodstream infections (18).

Interestingly, using the CLSI disk diffusion assay protocol (35), we found that FLC drug susceptibility and tolerance were significantly correlated for all three species, while BA drug susceptibility and tolerance were significantly correlated for *C. glabrata* and *C. parapsilosis*. This result differs from a previous screen of 219 *C. albicans* isolates that found no correlation between FLC susceptibility and tolerance when isolates were grown up on YPD and tested on casitone plates at 30 °C (16). Although the mechanisms of tolerance have not yet been well resolved, assay medium and temperature has previously been shown to significantly influence fungal drug susceptibility and can dramatically affect drug tolerance (15). The drivers and clinical importance of tolerance is an area of active interest and research.

The involvement of morphological changes in symptomatic vulvovaginal candidiasis is also an area of current interest. Hyphal formation is an important virulence trait in many infection contexts, and compounds that inhibit hyphal formation in *C. albicans* significantly reduce virulence in multiple contexts (36, 37). Our results show that *C. albicans* forms hyphae in the first 2 h regardless of FLC concentration, in alignment with previous research (26). Through the new quantitative platform we developed, we similarly found that percent area covered at the asymptote, biofilm growth rate, and time to reach the growth asymptote are also not correlated with FLC concentration, though we did uncover significant variation among isolates for these phenotypes. BA is clearly superior to FLC at the inhibition of hyphal formation, with the speed of morphological transition and subsequent biofilm formation influenced by BA in a dose-dependent manner. Hyphal formation was previously reported before to be responsible for vaginal inflammation (38, 39), and hyphae contribute to the invasive growth of *C. albicans* (20), suggesting that inhibiting the yeast to hyphal transition could help explain why BA is an effective treatment against complicated VVC.

If a biofilm has formed in the absence of a drug (e.g., prior to a symptomatic infection), penetration of drugs through the extracellular matrix is critical for eradication (40). We found that *C. albicans* biofilms are insensitive to FLC, consistent with past work (30, 31). Conversely, we found that BA can reduce biomass and activity of mature biofilms in a dose-dependent manner, and therefore is likely capable of penetration. Approximately 40% of the extracellular matrix of *C. albicans* is composed of polysaccharides and the major carbohydrates present is glucose (∼40%) (22, 41). Saccharides (especially glucose) have a strong affinity for borate (42), thus we think that BA likely penetrates the extracellular matrix of the biofilm by breaking the matrix apart. Compared to planktonic cells, *C. albicans* cells in a biofilm have a significantly higher β-1,3 glucan content in their cell wall, which has been implicated in FLC impermeability (43). We observed an increase in activity at high BA concentrations even though the biomass was decreasing. We think that this increase in activity is artificial and due to how the XTT activity assay works. Mitochondrial succinoxidase, cytochrome P450 systems, and flavoprotein oxidases convert XTT to a colored formazan (44). If high concentrations of BA degrade the extracellular matrix, then more XTT will get inside the cells, which is read out as increase in activity. Our future work will thus directly investigate the ability of BA to penetrate the cell wall of different *Candida* morphologies.

## Conclusion

Our work provides baseline data for multiple ways boric acid is an effective drug against diverse isolates of *Candida* species. We find that planktonic susceptibility and tolerance are more similar among isolates in boric acid than fluconazole. Boric acid is also effective at slowing or stopping the yeast-to-hyphal transition and biofilm growth in *C. albicans*, as well as able to penetrate and degrade mature biofilms. Planktonic responses in fluconazole and boric acid were not correlated, and nor did planktonic and biofilm phenotypes. Combined, this work advances our understanding of multiple biological fronts about why boric acid may be an effective treatment against vulvovaginal candidiasis.

## Material and Methods

### Isolates

235 *Candida* spp. isolates were used in this study (Supporting Materials Table 1). 155 of these isolates (85 *C. albicans*, 50 *C. glabrata*, and 20 *C. parapsilosis*) were provided by the clinical microbiology lab in the Health Sciences Centre (HSC) in Winnipeg, Canada. 80 additional *C. albicans* isolates that have previously been published from our lab freezer inventory were also examined.

**Table 1:**
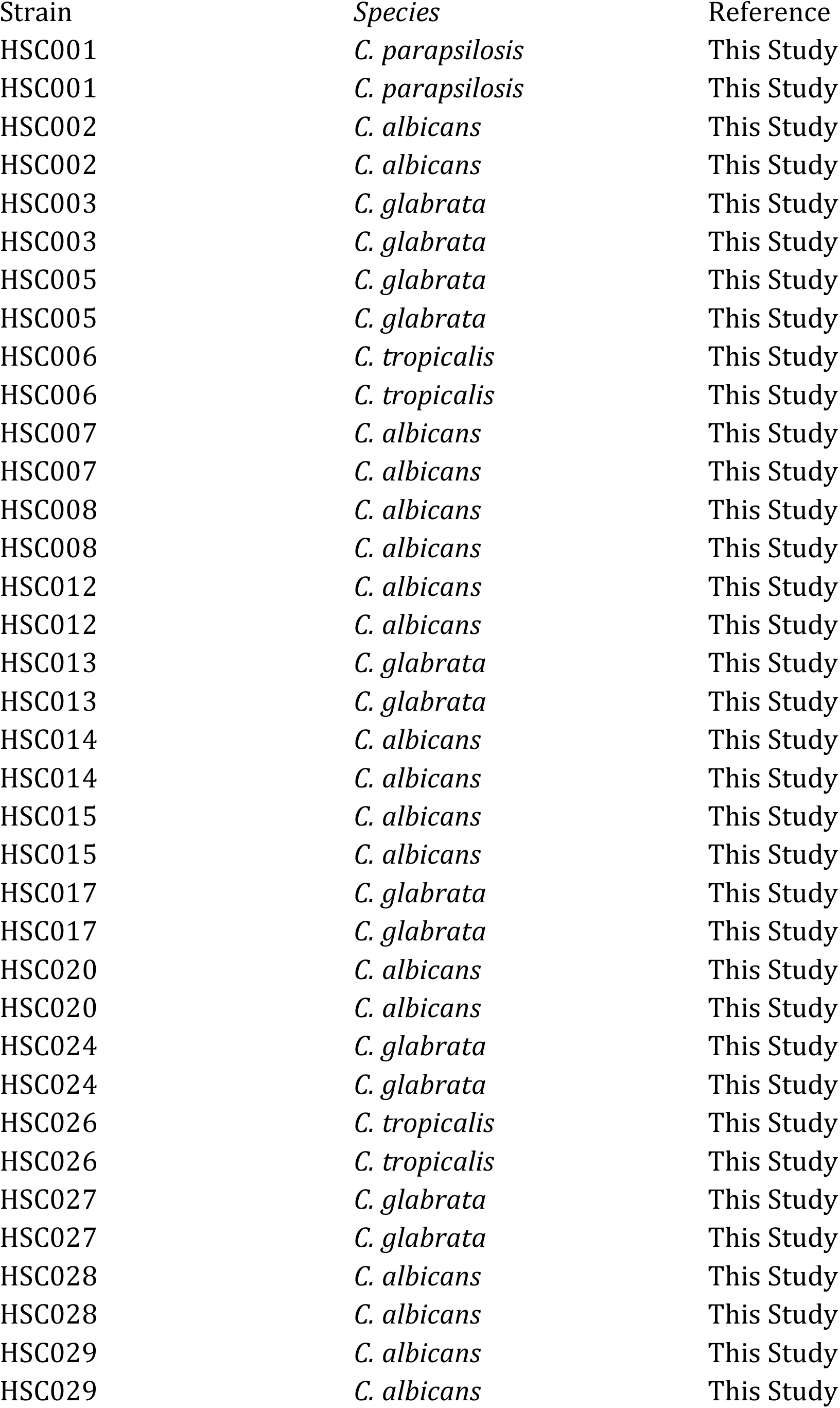
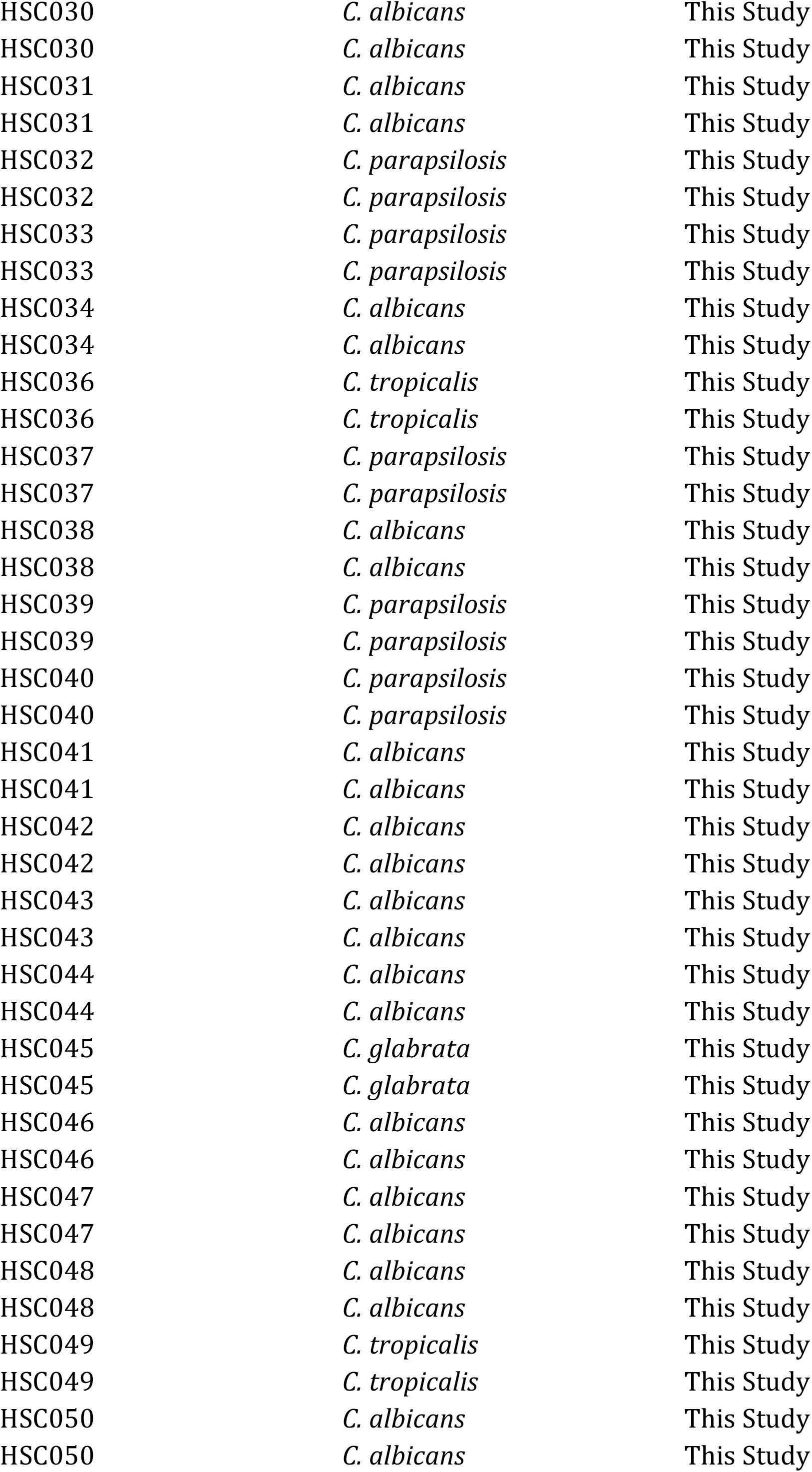
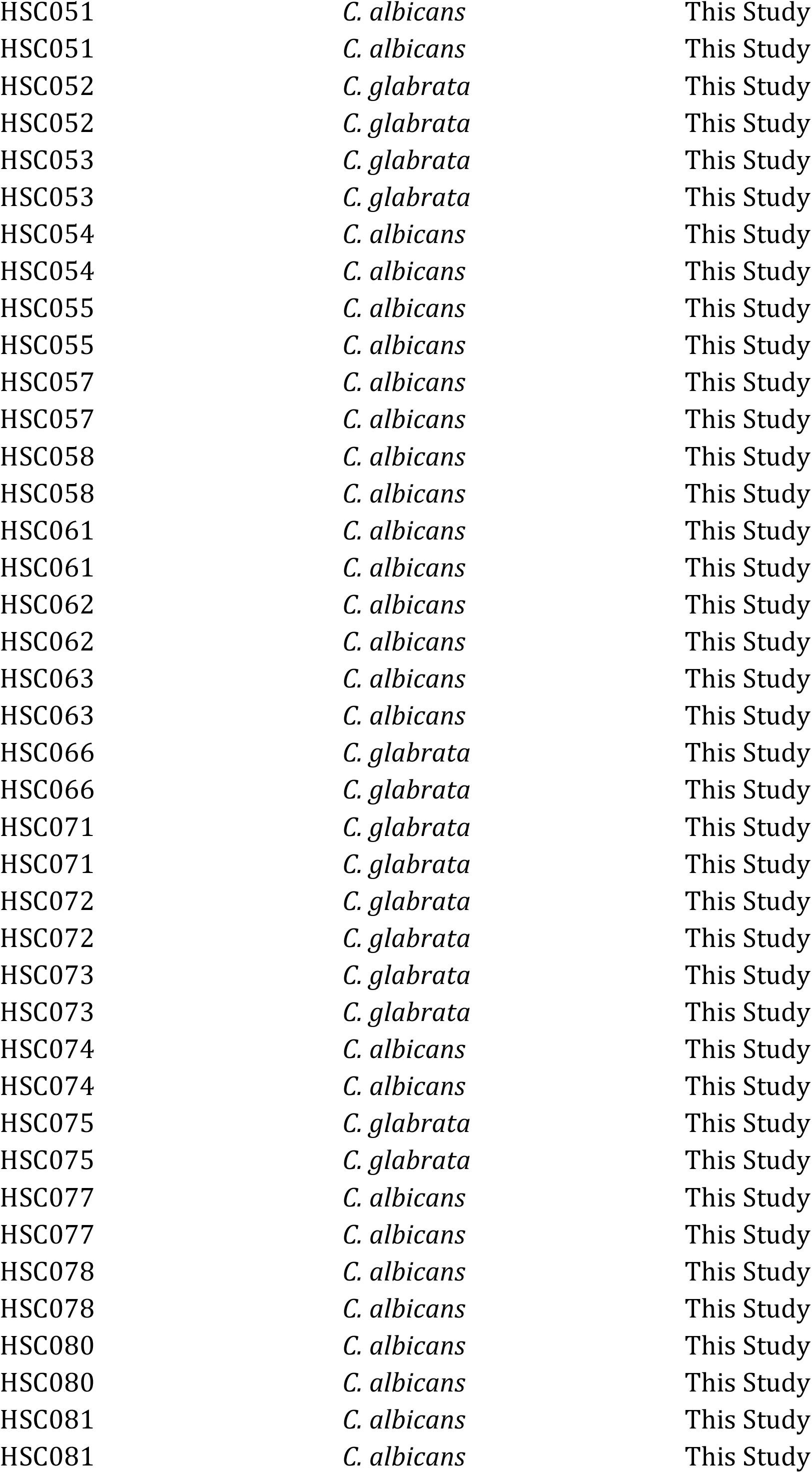
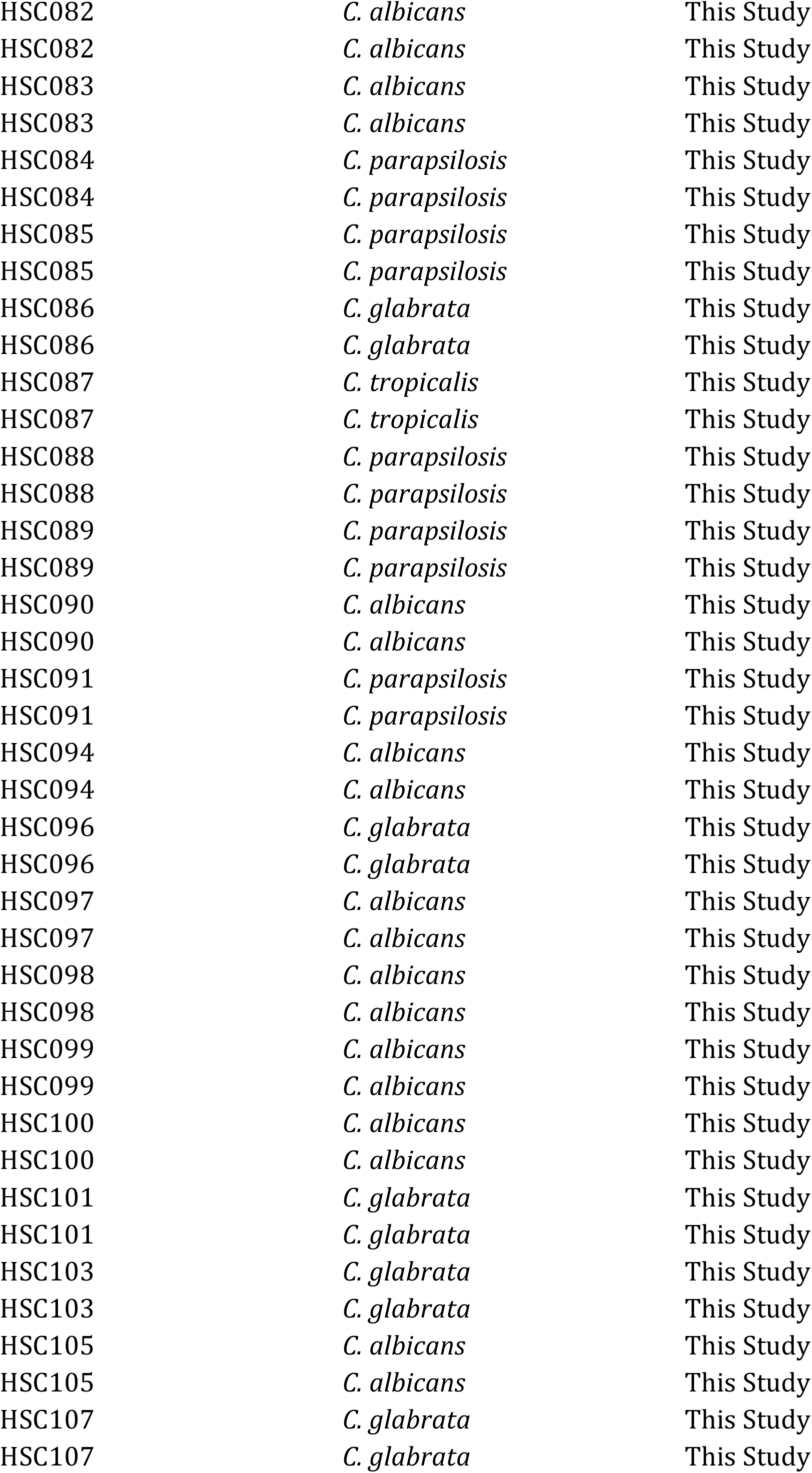
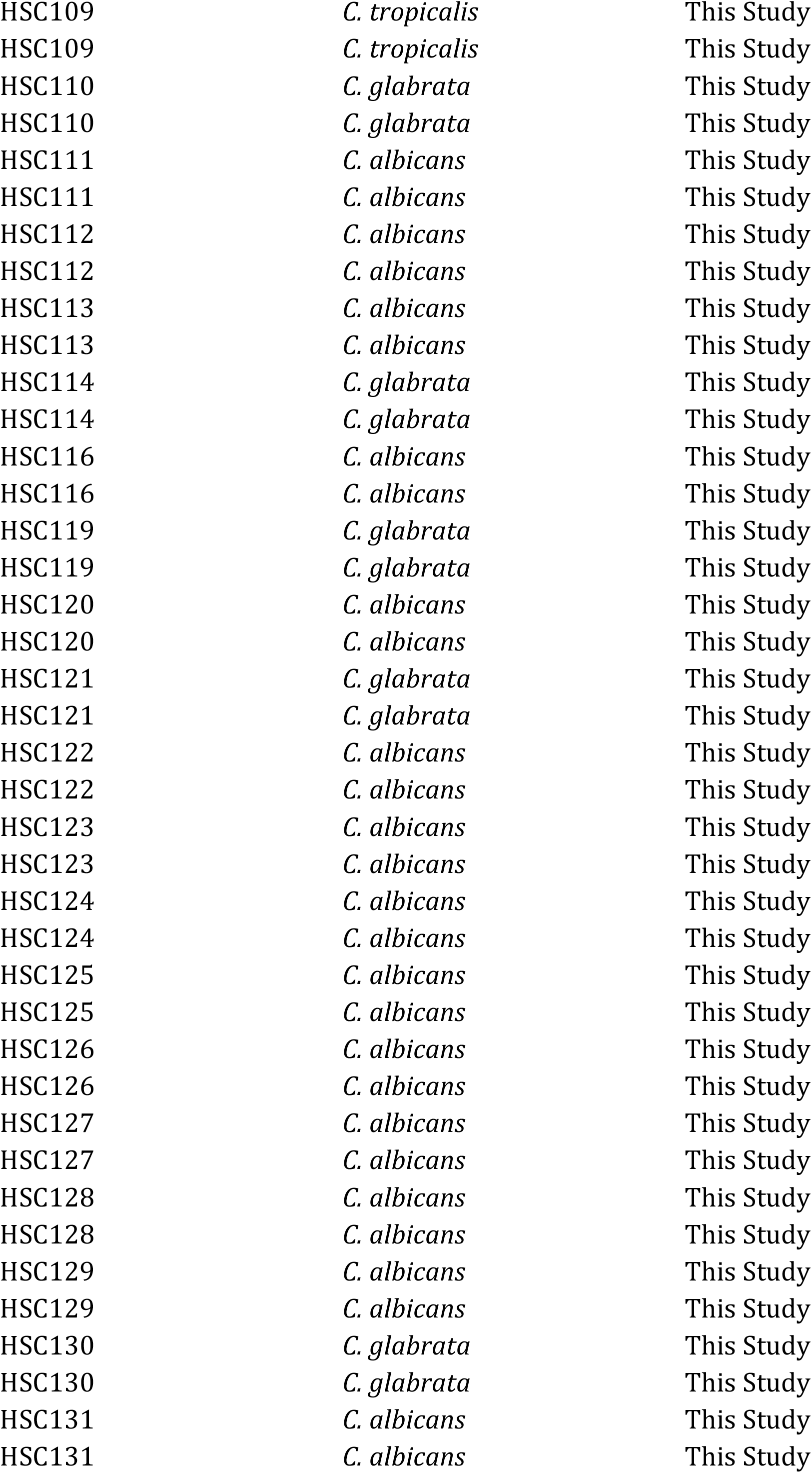
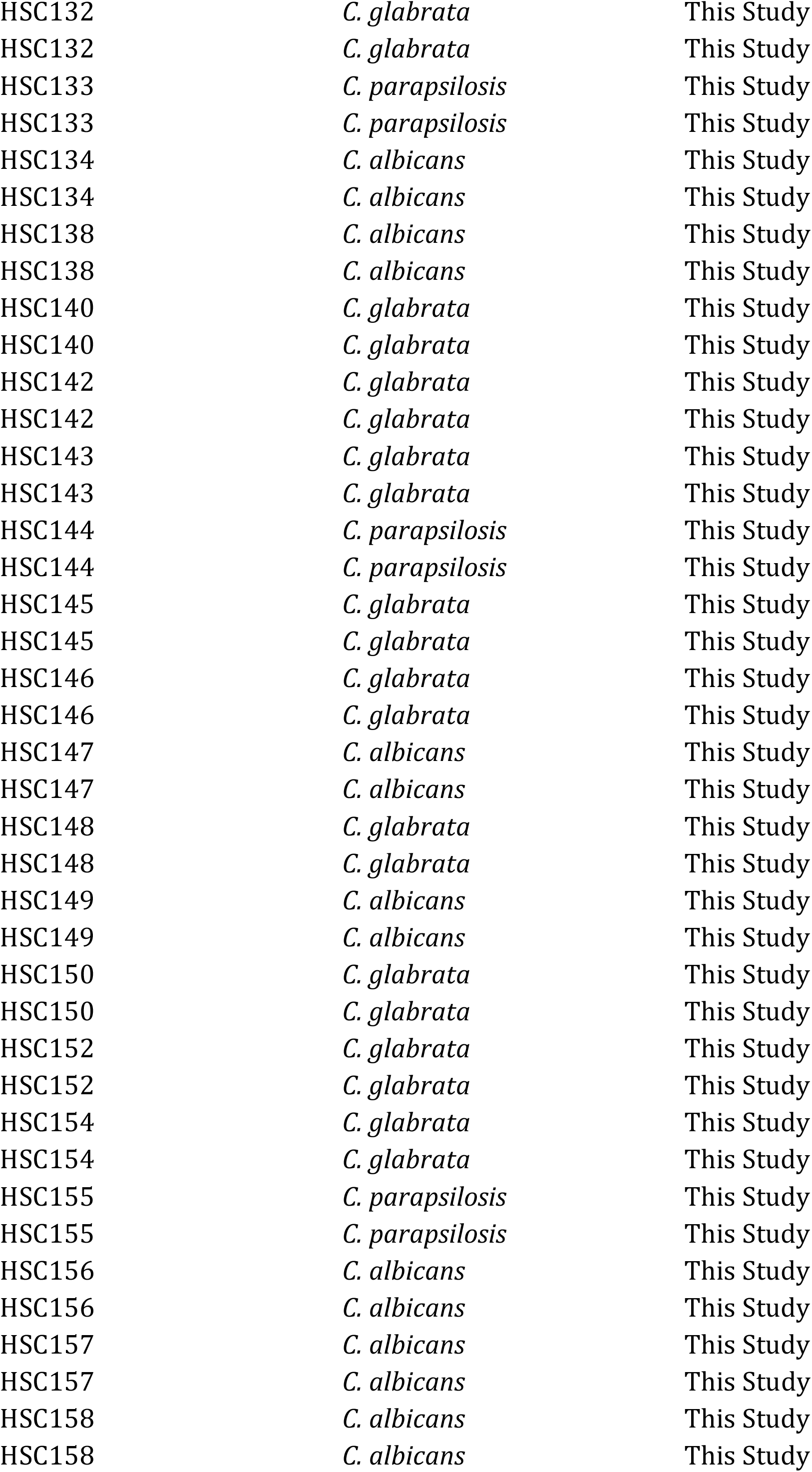
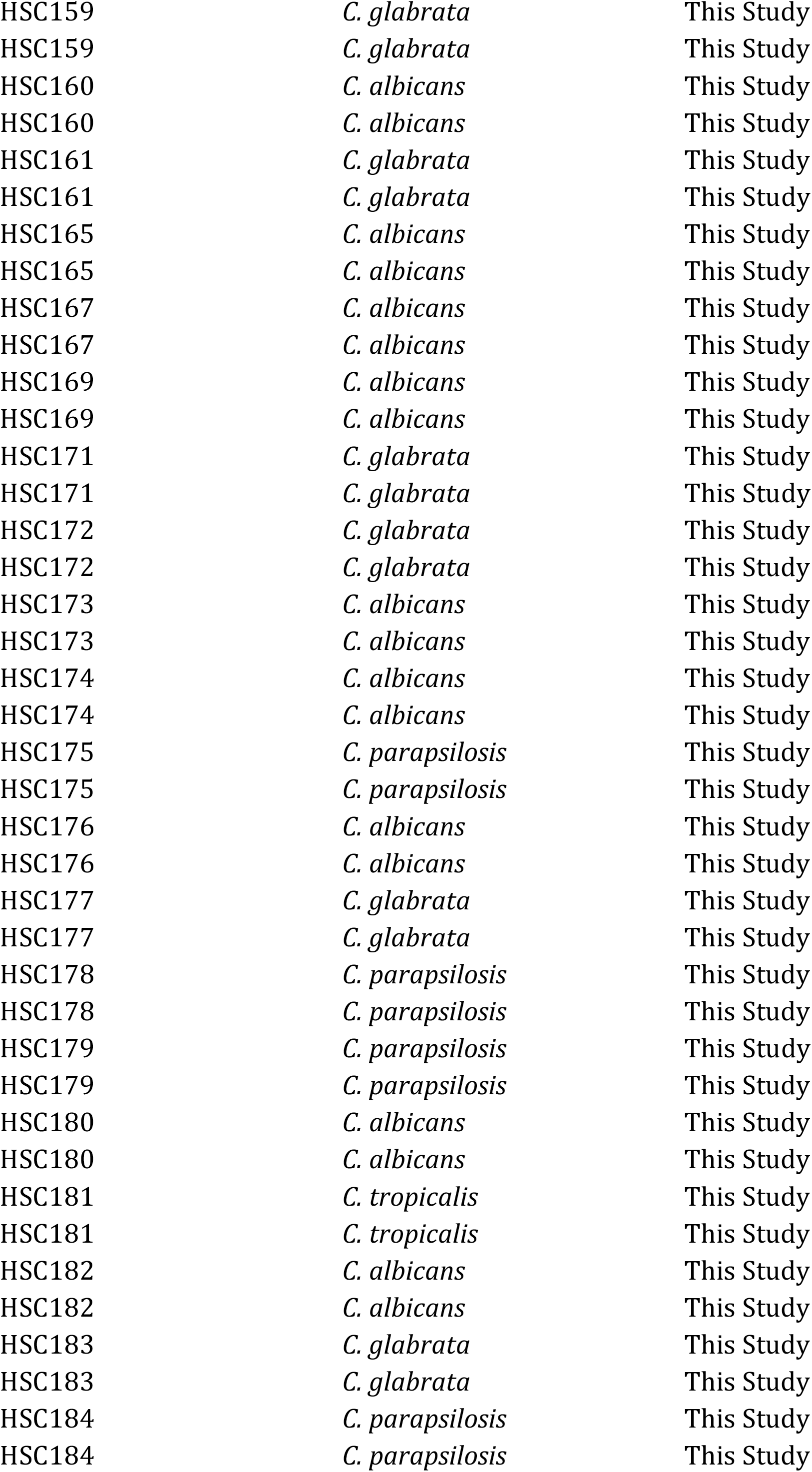
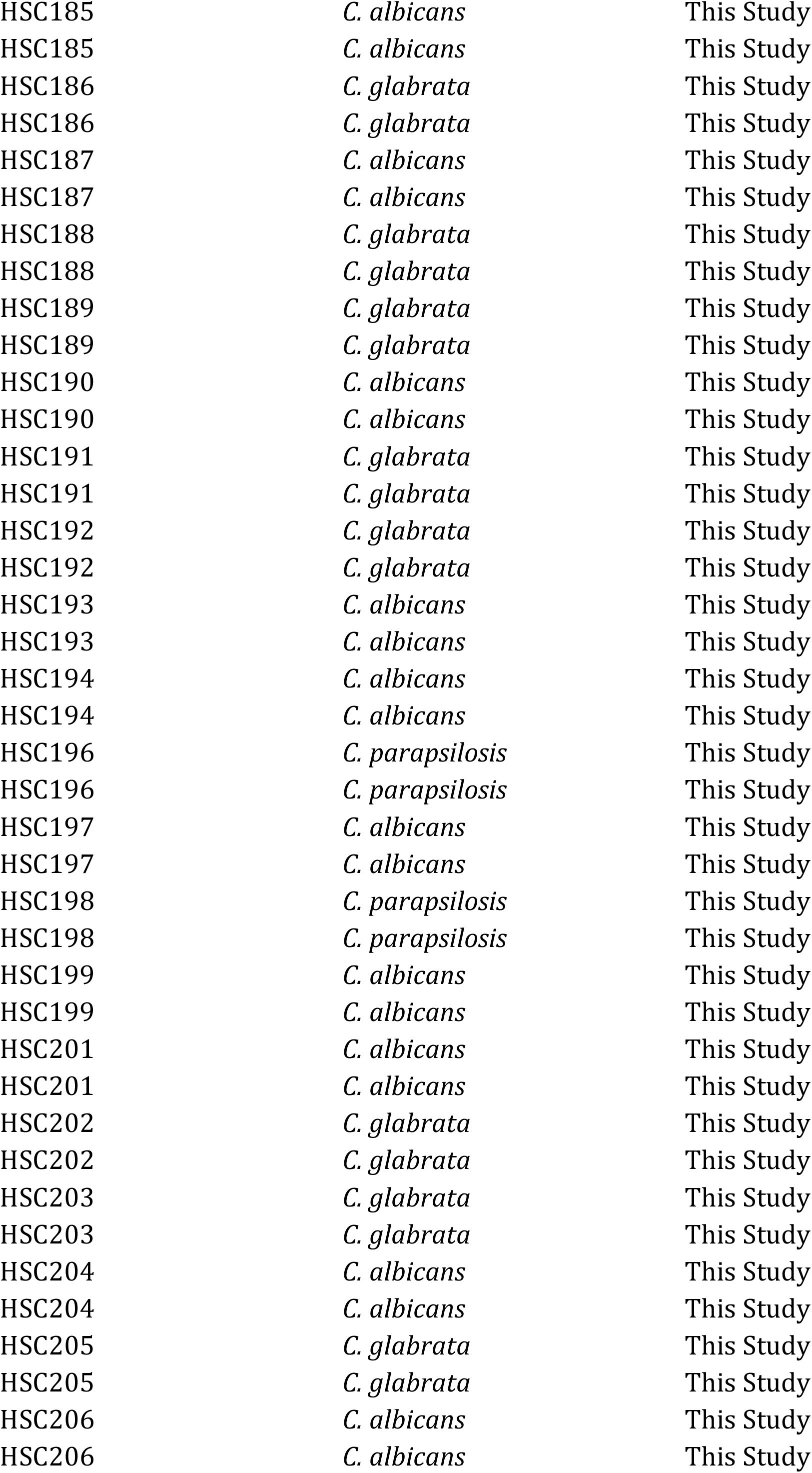
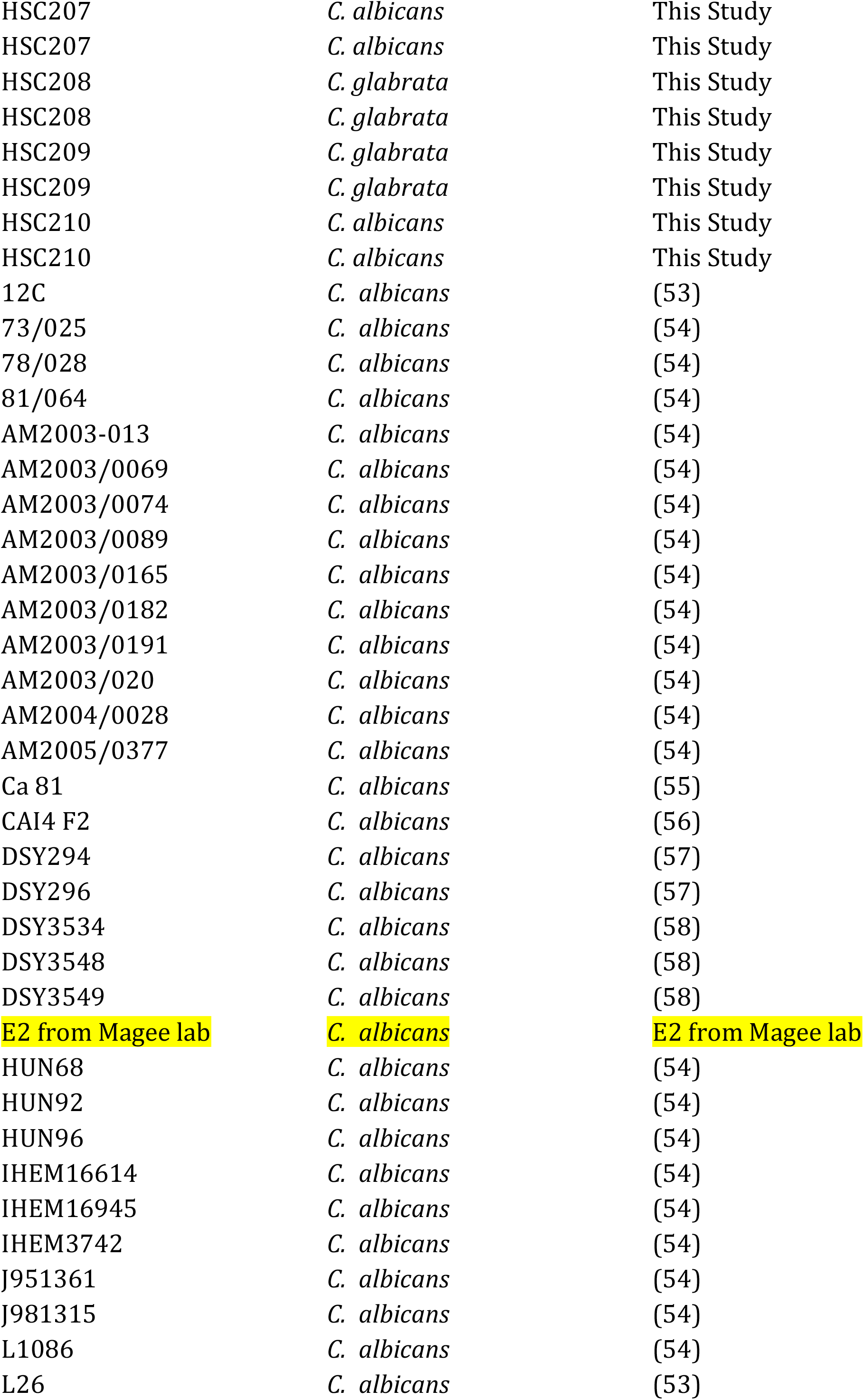
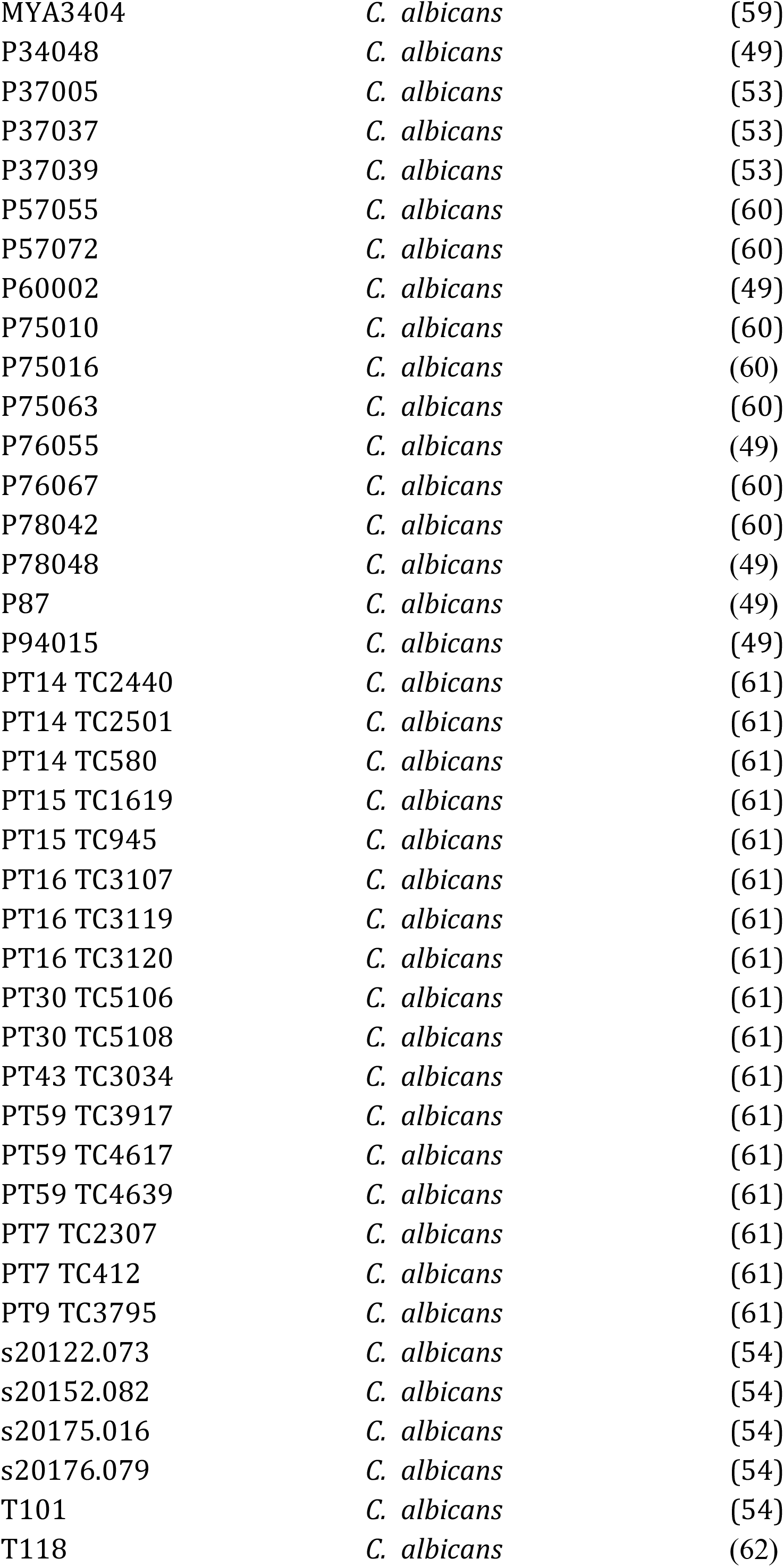
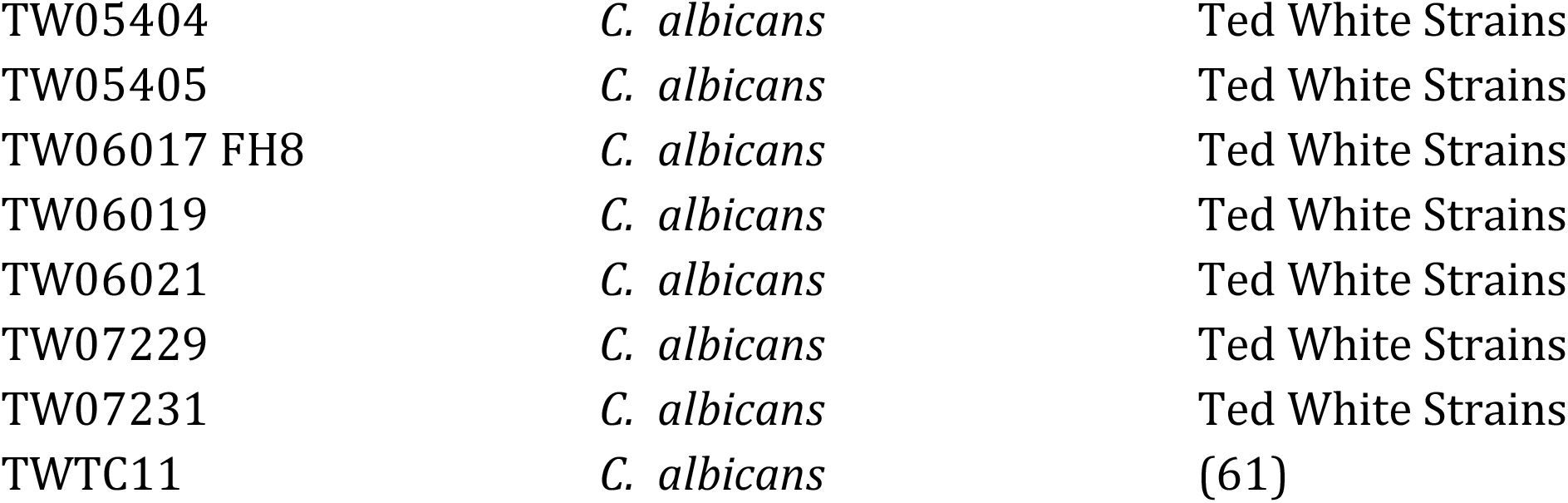
List of Isolates Used.

### Disk Diffusion Assay

To screen for drug susceptibility and tolerance, the CLSI document M44 guidelines for antifungal disk diffusion susceptibility testing were followed with slight modifications (35). Briefly, isolates were streaked from frozen glycerol stocks into Sabouraud Dextrose Agar (SDA) and incubated for 48 h at 37 °C. Colonies were resuspended in 200 μL of PBS and were standardized to an optical density (OD_600_) of 0.01 in 1 mL PBS. 100 μL of the standardized cells were spread in duplicates onto 15 mL Muller Hinton (MH) plates using sterile beads. 5 mg BA disks were prepared by transferring 10 μL of preheated 500 mg/mL BA in Dimethyl sulfoxide (DMSO) stock to blank antimicrobial disks (6 mm, Fisher Scientific). Single disks containing either 5 mg BA or 25 μg FLC (6 mm, Fisher Scientific) were placed in the center of the MH plates. Plates were then incubated at 37 °C for 48 h, and individual photographs were taken of each plate every 24 h and 48 h. Images were edited using ImageJ (45) according to recommendations specified in diskImageR vignette V2 146 (46). Disk diffusion analysis to measure drug susceptibility and drug tolerance was done using diskImageR package and the R script available at https://github.com/acgerstein/diskImageR/blob/master/inst/walkthrough.R (15). Susceptibility was measured from 48 h images as RAD_20_ (the radius where 20% growth of reduction occurs) while tolerance was measured as FoG_20_ (the fraction of growth above RAD_20_).

### Boric Acid levels

We picked the concentrations of BA based by measuring the minimum inhibitory concentration (MIC) for the isolates used in biofilm formation and eradication was determined using the CLSI M27-A2 guidelines with some modifications. Briefly, isolates used in biofilm formation and biofilm eradication screening were streaked from frozen stocks on SDA plates and were incubated overnight at 37 °C. Colonies were suspended in PBS and were standardized to an OD_600_ of 0.01 in 1 mL of RPMI-1640. 100 μL of the standardized culture were transferred to a 96-well plate (Greiner Bio-One) containing two-fold dilutions of BA with the maximum concentration of 50 mg/mL (maximum solubility of BA in RPMI) in column 1, column 11 containing no drug and column 12 containing just RPMI-1640. MIC_50_ were determined by taking OD_600_ after 24h and 48h. The MIC_50_ of all isolates was either 0.39 or 0.78 mg/mL and the MIC was 1.56 mg/mL. We decided to pick two levels above and below the MIC and to round the concentrations to one decimal place (i.e., 0.4, 0.8, 1.6, 3.2 and 6.4 mg/mL).

### Biofilm Formation Drug Response

The ability to form a biofilm in the presence of a drug of eight *C. albicans* isolates was measured as previously described in (47). Briefly, 100 μL of RPMI-1640 was added to all the wells of columns 1-9 and 12 of a flat-bottom 96-well microtiter plate (Greiner Bio-One). 200 μL of RPMI-1640 + 12.8 mg/mL BA was added to all wells of column 11 and 200 μL of RPMI-1640 + 64 μg/mL FLC was added to all wells of column 10. A 2 fold serial dilution for each drug was done so that odd-numbered columns (11-3) contain dilutions of BA, even-numbered columns (10-2) contain dilutions of FLC, and columns 1 and 12 contain no drug. *Candida albicans* isolates SC5314 (ATCC MYA-2876), FH1 (48), P87 (49), GC75 (49), P78048 (49), P75016 (49), P76055 (49) and T101 (50) isolates were grown overnight 30 °C in liquid YPD from frozen glycerol stocks. These isolates were picked because they span the phylogenetic diversity of *C. albicans*. Overnight cultures were washed with PBS and standardized to OD_600_ of 0.005 in 1.5 mL RPMI-1640. 100 μL of the standardized cultures were inoculated into the 96-well microtiter plate that contains our drugs. The plate was sealed with a clear seal (Thermo Scientific) and polystyrene microplate lid (Greiner Bio-One), and parafilm was used to seal the sides of the plate. The plate was incubated at 37 °C and images were taken manually every 1 h for 24 h using Evos FL Auto 2 inverted microscopy (Thermo Scientific).

### Microscopic Image Analysis

Time to form hyphae was recorded manually by going through the images and taking note of the time of the first hypha that was observed. Important parameters (% area at the asymptote, growth rate, time to reach the asymptote) that correlate with biofilm formation ability were extracted using a high throughput automated method (47). Briefly, Orbit Image Analysis (29) was trained on 14 images and a detection model with 99.3% correctly classified instances was created. This detection model was used to calculate the % area covered by cells for all of the images. Custom R scripts (47) were used to extract % area at the asymptote, growth rate, time to reach the asymptote from Orbit output.

### Biofilm Eradication

Biofilms were prepared as described previously with slight modifications (51). Briefly, overnight cultures of 68 different *C. albicans* isolates were prepared by inoculating 10 μL of frozen glycerol stocks in 500 μL YPD (Yeast Peptone Dextrose) liquid medium in a microplate shaker (about 200 rpm) at 30 °C overnight. Isolates were then standardized to OD_600_ of 0.01 in 1.5 mL RPMI-1640 broth and aliquots of 100 μL of the standardized culture were inoculated into flat-bottom 96-well microtiter plates (Fisher Scientific). The plates were then covered with a sealing membrane, sterilized lids, and the sides of the plates were sealed with parafilm. The plates were incubated statically at 37 °C for 24h. Mature biofilms were washed 3 times with phosphate buffer saline (PBS) to remove the non-adherent cells, and OD_600_ was taken to quantify the pre-drug biomass of the biofilms.

The examined drug concentrations ranged from 0.4 mg/mL to 12.8 mg/mL for BA and from 1 μg/mL to 32μg/mL for FLC. The biofilm antifungal susceptibility testing was done according to (51) with slight modifications. Briefly, the mature biofilms were treated with either BA or FLC and incubated statically with the drug for 24 h at 37 °C. The drug-treated biofilms were washed 3 times with PBS, and OD_600_ was taken to quantify the post-drug biomass of the biofilms. The biomass of the biofilms was normalized by subtracting the biomass of the biofilms pre-drug exposure from the biomass of the biofilms at each concentration. The metabolic activity was determined using the calorimetric 2,3-Bis-(2-Methoxy-4-Nitro-5-Sulfophenyl)-2*H*-Tetrazolium-5-Carboxanilide (XTT) assay, which measures mitochondria reduction of the tetrazolium salt reagent; briefly, mature biofilms were incubated with 100 uL of XTT for 2 h in 37 °C, and OD_490_ was taken to quantify the metabolic activity. Biofilm activity was normalized by subtracting the activity at no drug concentration from the activity at each drug concentration. We excluded 4 isolates from our analysis because they failed to form biofilms during the pre-drug exposure phase.

### Statistical Analysis

Values for susceptibility and tolerance screening represent the average of three biological replicates with one or more technical replicates each. Values for hyphal and biofilm formation represent the average of two biological replicates with one technical replicate each. Values for biofilm eradication represent the average of three biological replicates with one technical replicate each. Error bars throughout represent standard error.

All statistical tests performed and graphs generated were done using R programming language (52). Since our data is not normally distributed, we used rank-based measures. When comparing susceptibility and tolerance among species for each drug, we used Kruska-Wallis rank-sum test and determined significance using post-hoc pairwise Wilcoxon tests with the (Benjamini and Hochberg, 1995) *P*. We looked at homogeneity of group variances using Flinger-Killeen test of homogeneity. We used Spearman’s rank correlation when looking at the correlation between planktonic drug responses. We used linear mixed-effect models from lmerTest R package (32) to determine if planktonic drug responses would influence biofilm drug responses. In all cases, the specific statistical test is indicated inline. Significance was assigned for *P* <0.05. Raw data and scripts to generate figures and statistical analyses are available at https://https://github.com/acgerstein/BAFLC_Phenotypic

## Acknowledgements

We thank Dr. Markus Stein at the Health Sciences Centre in Winnipeg for providing isolates from the clinical microbiology lab, as well as Drs. Judith Berman, Ted White, Donna MacCallum, Dominique Sanglard, Peter Magee, Shawn Lockhart, Claude Pujol, David Soll for sharing strains with the Candida community. We are grateful to Dr. Vanessa Poliquin for her critical insight and thoughtful conversations about her experience treating women with complicated vulvovaginal candidiasis. We thank R. Kukurudz and K. Chokkar for laboratory assistance, and S. Harris, P. Pelka, S. Cardona and A. Kumar for use of equipment.

## Supplementary Materials

**Supplementary Figure 1.**
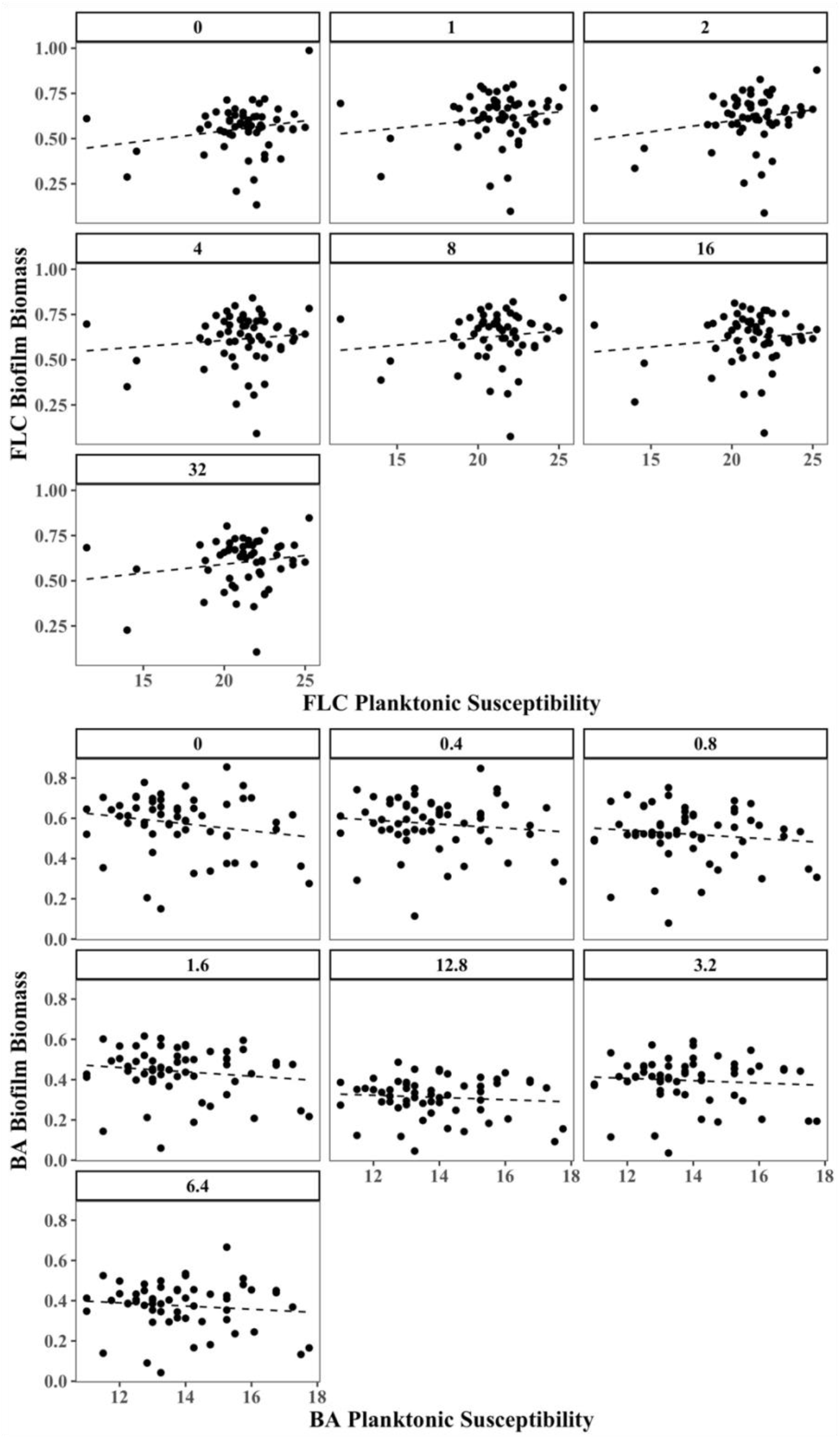
Association of planktonic susceptibility and biofilm biomass of *C. albicans* isolates. There is no association between planktonic susceptibility and biomass of the biofilms for both FLC and BA (linear mixed-effect model).

**Supplementary Figure 2.**
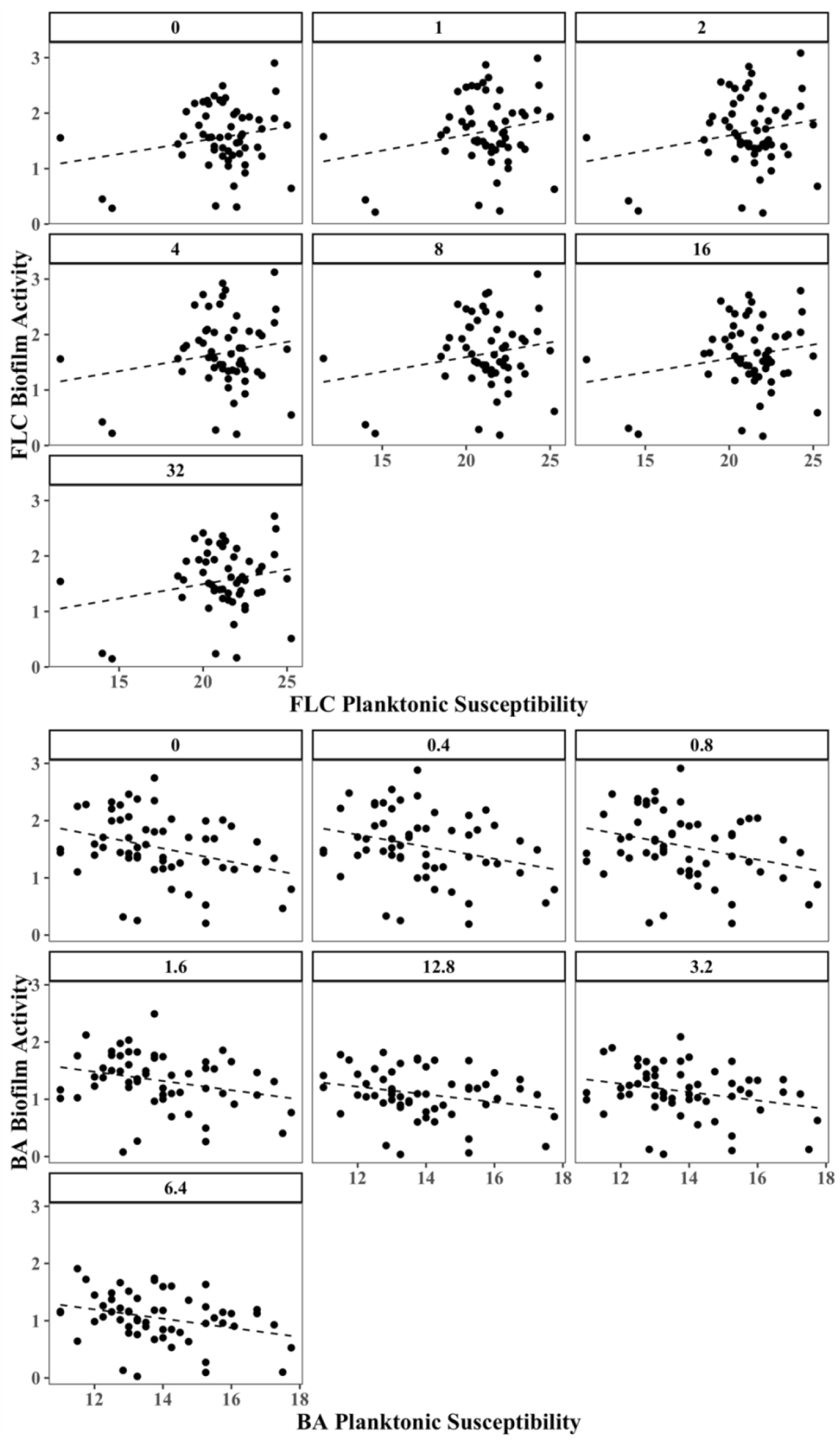
Association of planktonic susceptibility and biofilm activity of *C. albicans* isolates. There is an association between BA planktonic susceptibility and biomass of the biofilms; however, this association absent in FLC (linear mixed-effect model).

**Supplementary Figure 3.**
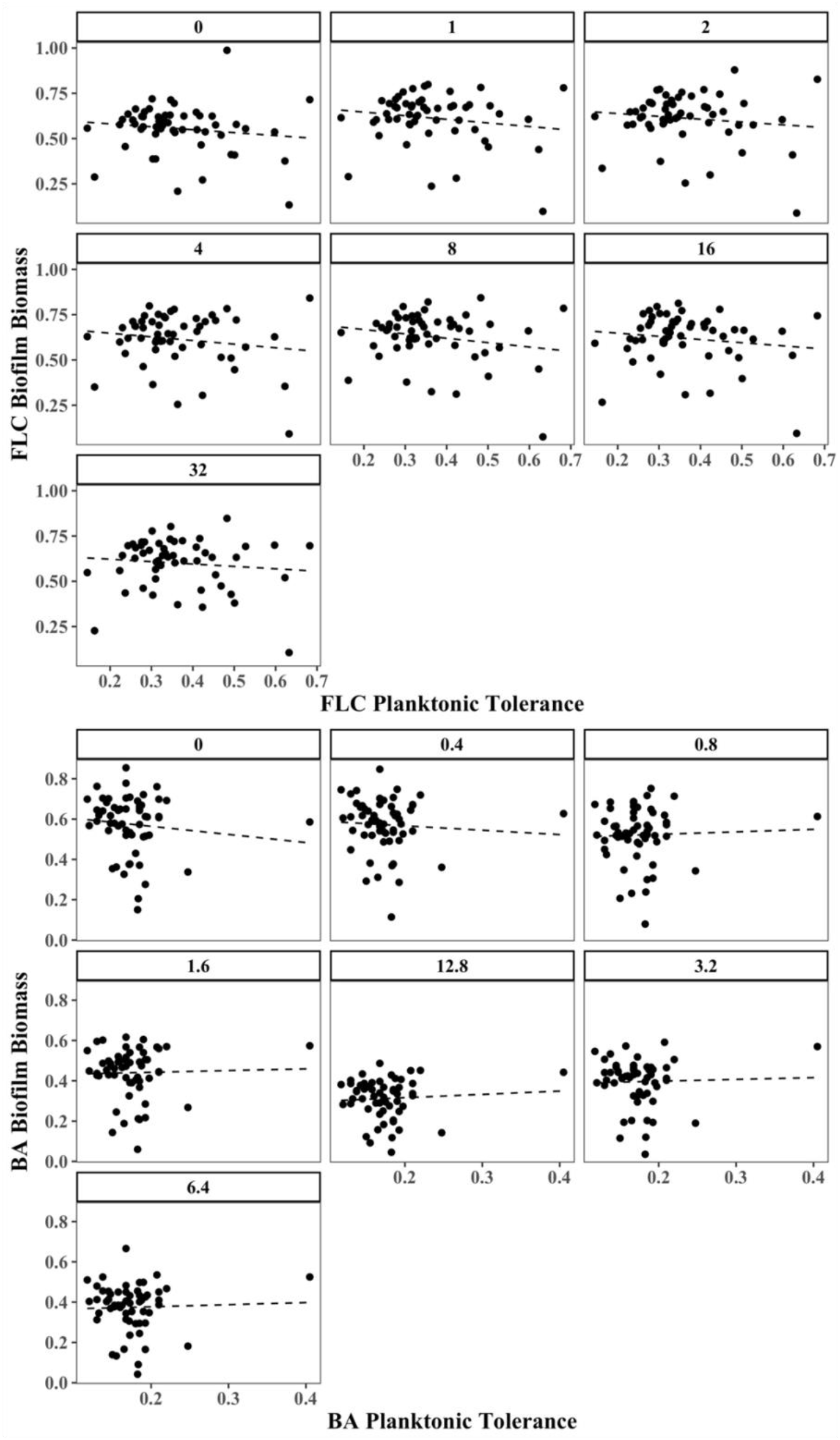
Association of planktonic tolerance and biofilm biomass of *C. albicans* isolates. There is no association between planktonic tolerance and biomass of the biofilms for both FLC and BA (linear mixed-effect model).

**Supplementary Figure 4.**
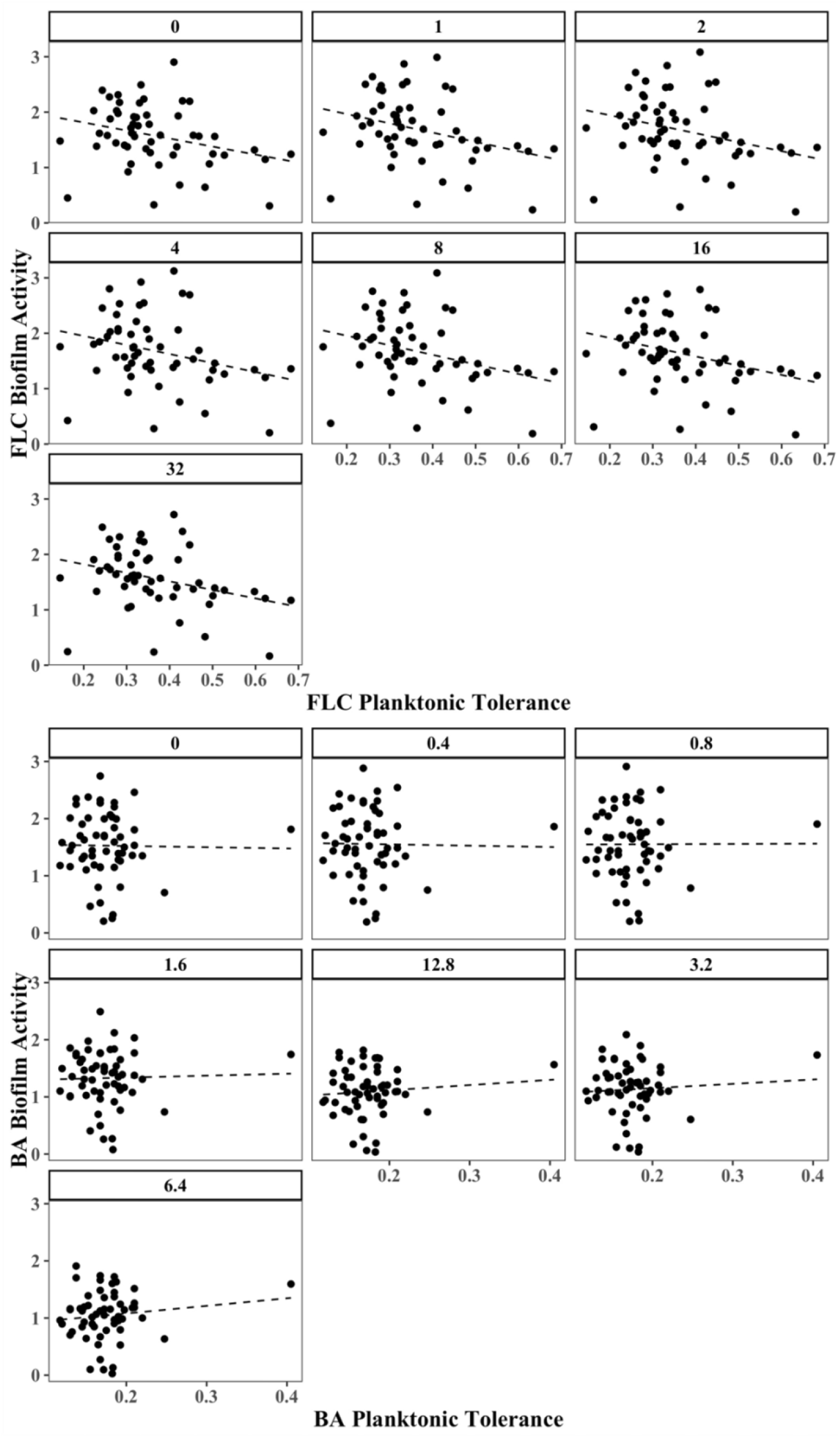
Association of planktonic tolerance and biofilm activity of *C. albicans* isolates. There is an association between FLC planktonic tolerance and activity of the biofilms (linear mixed-effect model); however, this association is absent in BA.

## Notes

https://github.com/acgerstein/BAFLC_Phenotypic

### Competing Interest Statement

The authors have declared no competing interest.

